# H_2_ consumption by various acetogenic bacteria follows first-order kinetics up to H_2_ saturation

**DOI:** 10.1101/2024.05.08.593002

**Authors:** Laura Muñoz-Duarte, Susmit Chakraborty, Louise Vinther Grøn, Maria Florencia Bambace, Jacopo Catalano, Jo Philips

## Abstract

Acetogenic bacteria play an important role in various biotechnological processes, because of their chemolithoautotrophic metabolism converting carbon dioxide with molecular hydrogen (H_2_) as electron donor into acetate. As the main factor limiting acetogenesis is often H_2_, insights into the H_2_ consumption kinetics of acetogens is required to assess their potential in biotechnological processes. In this study, initial H_2_ consumption rates at a range of different initial H_2_ concentrations were measured for three different acetogens. Interestingly, for all three strains, H_2_ consumption was found to follow first-order kinetics, i.e. the H_2_ consumption rate increased linearly with the dissolved H_2_ concentration, up to almost saturated H_2_ levels (600 µM). This is in contrast with Monod kinetics and low half-saturation concentrations, which have commonly been assumed for acetogens. The obtained biomass specific first-order rate coefficients (*k*_1_^X^) were further validated by comparison with values obtained by fitting first-order kinetics on previous time-course experimental results. The latter method was also used to determine the *k*_1_^X^ value of five additional acetogens strains. Biomass specific first-order rate coefficients were found to vary up to six-fold, with the highest *k*_1_^X^ for *Acetobacterium wieringae* and the lowest for *Sporomusa sphaeroides*. Overall, our results demonstrate the importance of the dissolved H_2_ concentration to understand the rate of acetogenesis in biotechnological systems.

## Introduction

Acetogens are an interesting group of strictly anaerobic, chemolithoautotrophic bacteria. They are phylogenetically diverse, but have the common characteristic of utilizing the Wood–Ljungdahl pathway (WLP) for CO_2_ fixation and acetate production while conserving energy, using H_2_ as electron donor (Basen & Müller, 2023), according to the overall reaction:

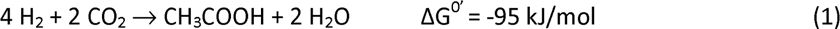

Acetogens are widely distributed across diverse environments, ranging from soils to the human gut (Drake et al., 2002; Smith, Shorten, Altermann, Roy, & McNabb, 2020). In addition, the metabolic abilities of acetogens are attractive for the development of novel biotechnological processes to convert carbon dioxide into valuable products. In gas fermentation, CO_2_ is metabolized in a bioreactor fed with H_2_ gas, while an electrode is used to produce H_2_ *in situ* in the case of microbial electrosynthesis (Harnisch et al., 2024). In these biotechnological settings, as well as in natural environments, the supply of H_2_ often limits acetogenesis (Philips, 2020). Understanding the relationship between the H_2_ level and the rate of H_2_ consumption and acetogenesis, i.e. H_2_ consumption kinetics, is thus essential for assessing the potential of acetogens in biotechnological and environmental processes.

Most often, H_2_ consumption and growth by acetogens is assumed to follow Monod kinetics (similar to Michaelis Menten kinetics for enzymes) (Philips, 2020), in which the substrate consumption rate (*r*) increases with the substrate concentration *[S]*, until a maximum is reached (Rittman & McCarty, 2020).

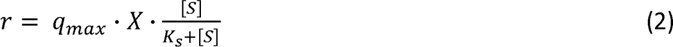

The half-saturation coefficient *K*_s_ gives the substrate concentration at which half of the maximum rate is reached. This maximum rate depends on the biomass-specific maximum consumption rate (*q*_max_) and the biomass concentration (*X*). So far, (maximum) growth rates or doubling times (from which *q*_max_ can be calculated using the growth yield) have frequently been reported for acetogens (Bengelsdorf et al., 2018; Groher & Weuster-Botz, 2016a), but only few studies have reported half-saturation coefficients for H_2_ consumption by acetogens (**Table 1**). Moreover, not all these studies carefully characterized whether the relationship between the H_2_ consumption rate and the H_2_ concentration is well described by Monod kinetics (Eq. 2).

**Table 1.** Reported values of half-saturation coefficients (*K*_s_) for H_2_ consumption by acetogens.

| Acetogen | $K_s$ ($\mu\text{M}$) | Method | Reference |
| --- | --- | --- | --- |
| *Acetobacterium woodii* | 0.73 | Chemostat; based on half of the maximum growth rate | (Peters et al., 1998) |
| *Acetobacterium woodii* | 2500 | Enzyme assay on purified hydrogenases from cells grown on fructose | (Ragsdale & Ljungdahl, 1984) |
| *Acetobacterium bakii* | 4 | Batch; H<sub>2</sub> depletion curves over time, constant biomass | (Kotsyurbenko et al., 2001) |
| *Sporomusa termitida* | 6 | Batch without headspace; H<sub>2</sub> depletion in the liquid | (Breznak et al., 1988) |
| *Sporomusa sphaeroides* | 341 | Enzyme assay on purified hydrogenase from betaine grown cells | (Dobrindt & Ulrich, 1996) |
| *Eubacterium limosum* | 340 | Batch; H<sub>2</sub> depletion curves over time- initial rates (similar as our procedure) | (Genthner & Bryant, 1987) |
| *Clostridium* strain P11 | 233 | Hydrogenase assay on permeabilized whole cells grown on syngas | (Skidmore et al., 2013) |
| *Clostridium ljungdahlii* | 327 | Hydrogenase assay on permeabilized whole cells grown on syngas | (Skidmore et al., 2013) |

Although acetogens perform the same overall reaction (Reaction 1), they are metabolically diverse, which is reflected by differences in their energy conservation strategies (Katsyv & Müller, 2020; Kremp et al., 2020, 2022; Rosenbaum & Müller, 2021; Wiechmann et al., 2020; Wiechmann & Müller, 2021), leading to different H_2_ thresholds, i.e. the H_2_ partial pressure at which acetogenesis halts (Munoz & Philips, 2023). H_2_ consumption kinetics likely also differ significantly among acetogenic strains, as different doubling times or growth rates have been reported (Groher & Weuster-Botz, 2016a; Philips, 2020). However, the few studies that have published half-saturation coefficients for H_2_ consumption by acetogens (**Table 1**), tested each just one acetogenic strain. Moreover, these studies used various approaches to quantify the kinetic parameters (**Table 1**), resulting in parameter values that are hard to compare among the different studies.

With this work, we aim to investigate how the H_2_ consumption rate of acetogens depends on the H_2_ concentration, as well as to compare the H_2_ consumption kinetics of different acetogens. We first measured initial H_2_ consumption rates at different initial H_2_ levels (ranging from ∼0.5 to 600 µM) for three different strains under the same conditions. These analyses showed that H_2_ consumption follows first-order kinetics, rather than Monod kinetics, as no saturation was observed over the whole range of tested H_2_ levels. The obtained first-order rate coefficients were further validated by comparison with values fitted on previous time-course experimental results (Munoz & Philips, 2023). Moreover, first-order kinetic coefficients of five additional acetogens strains were also obtained by fitting time-course experimental results.

## Materials and methods

### Strains and culture conditions

The acetogenic strains *Sporomusa ovata* (DSM 2662), *Acetobacterium woodii* (DSM 1030) and *Clostridium ljungdahlii* (DSM 13528) were revived from frozen stocks in the recommended medium (DSMZ 311, 135 and 879, respectively) using an organic substrate (betaine or fructose). After one transfer in the same medium, when an optical density (OD_600_) of approximately 0.6 was reached, the cultures were filtered through a 0.2 µm polyethersulfone membrane. The cells were washed three times with autotrophic growth medium inside an anaerobic chamber with an N_2_/CO_2_ atmosphere (Jacomex, France) to remove acetate and organic substrates resulting from the heterotrophic growth medium. The washed cells were resuspended in autotrophic growth medium, with a composition of 1 g NH_4_Cl, 0.1 g KCl, 0.8 g NaCl, 0.1 g KH_2_PO_4_, 0.077 g MgCl_2_, 0.02 g CaCl_2_.2H_2_O, 0.3 g L-Cysteine-HCl.H_2_O, 20 mM NaHCO_3_, 50 mM 3-(N-morpholino) propane sulfonic acid (MOPS) brought to pH 7.0 with NaOH, 0.1 g yeast extract, 10 mL vitamin solution DSMZ 141, 1 mL trace element solution and 0.1 mL tungstate-selenite solution per liter (Patil et al., 2015). The strains were not grown on H_2_ before the start of the experiment, to avoid possible adaptations to high H_2_ levels (Kato et al., 2015).

### Initial consumption rate experiment

The H_2_ consumption of the three acetogenic strains was characterized at a range of different initial H_2_ concentrations, in the same conditions of temperature, medium composition, electron acceptor concentration, pH (7) and agitation. Serum bottles (120 mL) were filled with 40 mL of autotrophic growth medium, while the headspace was flushed with gas mixtures with different H_2_ concentrations (ranging from ∼0.1 to 60%) and with 20% CO_2_ in N_2_. These gas mixtures were prepared using Brooks SLA5800 mass flow controllers. The serum bottles were pressurized to 1.2 bar absolute pressure. At least five different H_2_ initial concentrations were tested in triplicate for each strain. The gas phase was allowed to equilibrate with the liquid phase by agitation on an orbital shaker (Gerhardt, DE) at 100 rpm overnight at 30 °C. After equilibration, the bottles were inoculated with 5 mL of washed-cell suspension to reach a final starting OD_600_ of 0.2. The bottles were incubated at 30 °C and shaken at 100 rpm, while kept upside down to avoid gas loss through the rubber septa.

After inoculation, the cells were given at least 1 hour of equilibration time. After that period (set as time zero), samples (1.5 mL) were taken from the headspace for GC analysis until the H_2_ concentration had decreased by 10% or more of the initial H_2_ concentration. The total pressure was measured using a digital manometer (Omega DPG108, UK) before each headspace sampling. This manometer was connected to a needle (25G x 5/8”) using a gas-tight 3D-printed adapter, minimizing the volume (250 µL) sampled for each pressure measurement. The partial pressure of H_2_ was calculated from the headspace H_2_ concentration (%) measured by GC and the total pressure at each sampling point. The concentration of dissolved H_2_ in the liquid phase was calculated using the Henry constant for H_2_ (7.8·10^−4^ M bar^−1^, at 30 °C) (Sander, 2023). Abiotic controls with different initial H_2_ concentrations were sampled using an identical protocol to confirm that the H_2_ depletion rates resulted from the metabolic activities of the acetogens. Liquid samples were taken for measurement of the acetate concentration in the initial and final sampling time. A parallel experiment with the same initial H_2_ concentrations was sampled over time to monitor the change in OD_600_.

### Analytical methods

The analytical methods to measure the H_2_ headspace concentration by GC, acetate concentration by GC-VFA and the OD_600_ by spectrophotometry were previously described (Munoz & Philips, 2023). The headspace H_2_ concentration was analyzed using a CompactGC 4.0 (Interscience, Netherlands). This gas chromatographer has a Thermal Conductivity Detector (TCD) operated at 90°C and two columns in line: Molsieve 5A 30 m x 0.32 mm and Rt-QBond 3 m x 0.32 mm with argon as carrier gas at 40°C. The software is Chromeleon 7. The CompactGC was calibrated with H_2_-containing gas mixtures prepared by gas dilutions with Brooks SLA5800 mass flow controllers.

In addition, correlation curves of the cell density and dry weight (DW) with the optical density (OD_600_) were obtained for each acetogenic strain (**Table S1**). The cell density (cells mL^−1^) was quantified by Impedance Flow Cytometry (IFC) (Bactobox, SBT, Denmark) following the manufacturer’s protocol. A heterotrophically grown culture was washed and resuspended in autotrophic growth medium as described above, and thoroughly homogenized using a vortex. Afterwards, culture samples of about 100 - 500 µL were diluted in 10 mL of Phosphate Buffer Saline (pH 7.4 NaCl (13.7 mM), KCl (0.27 mM), Na_2_HPO_4_ (1.01 mM), KH_2_PO_4_ (0.176 mM)) with a conductivity between 1900 - 2100 µS/cm and measured in triplicates with IFC. Total cell numbers were used to calculate the correlation factors. The IFC method was validated using qPCR on the bacterial 16S rRNA gene (**Supplementary Method S1**), which gave results in the same order of magnitude. Likewise, the dry weight was correlated to OD_600_ for each acetogenic strain in quintuplicates. Cell suspensions were dried in an oven at 80°C until constant weight was reached and the average dry weight was used to calculate the correlation factors.

### Determination of kinetic order and kinetic coefficients from the initial rate experiment

The rate of H_2_ consumption was correlated to the initial H_2_ concentration to determine the kinetic order type. The rate of H_2_ consumption was derived from the decrease of the H_2_ partial pressure over time. Only the initial data points, corresponding to about 10% change of the initial H_2_ concentration, were used to calculate the H_2_ consumption rate at an almost constant H_2_ concentration. The small loss of H_2_ related to the withdrawal of the gaseous samples was estimated by measuring the total pressure before and after sampling in control bottles. This loss of H_2_ due to sampling was subtracted from the overall decrease of the total mole H_2_ (Dopffel et al., 2023) to avoid overestimating the microbial consumption. The volumes of gas and liquid phases were considered constant throughout the experiment. Our calculations assume that H_2_ was the only limiting substrate and that the agitation resulted in instantaneous equilibrium between the gas and liquid phases in the bottles. The latter assumption is further evaluated and discussed below.

As explained in the result section below, H_2_ consumption followed first-order kinetics over the range of H_2_ concentrations evaluated in this study. This entails that the H_2_ consumption rate *r* increases linearly with the dissolved H_2_ concentration *c*_H2_. The slope of this linear increase is given by the first order rate coefficient *k*_1_, which can also be written as the biomass-specific first-order rate coefficient 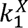 and the biomass concentration X:

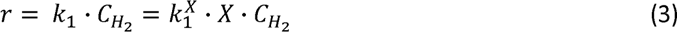

Equation (3) derives from Monod kinetics (Equation 2), in case the H_2_ concentration is much lower than *K*_s_.

Equation (3) can further be extended to also account for the H_2_ threshold 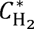, which is the minimum dissolved H_2_ concentration at which acetogenesis is thermodynamically feasible, and below which acetogenesis becomes unfavorable and halts (Munoz & Philips, 2023). Incorporating the H_2_ threshold as previously proposed (Kotsyurbenko et al., 2001; Smith, Shorten, Altermann, Roy, & Mcnabb, 2020), gives:

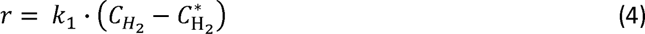

In our experimental setup, the H_2_ consumption by microbial metabolism only occurred in the liquid phase, but dissolved H_2_ was replenished by the dissolution of gaseous H_2_. For this reason, the first-order kinetic coefficients were determined based on the total mole change over time, similar as described previously (Karadagli & Rittmann, 2007). The total moles *(n*_T,H2_) of H_2_ at any time point is described by Eq. 5:

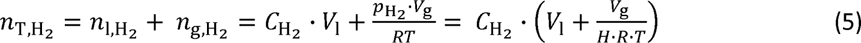

With *n*_J_ moles in the liquid phase, *n*_g_ moles in the gas phase, *v*_J_ the volume of the liquid phase, *v*_g_ the volume in the gas phase, *µ*_H2_ the partial pressure of H_2_ in the headspace, *R* the ideal gas constant, *T* the temperature, *H* Henry’s constant of H_2_ at 30 °C. Combining Equations (4) and (5), gives the ordinary differential equation describing the change of the total H_2_ moles in the system over time, depending on microbial H_2_ consumption considering first-order kinetics :

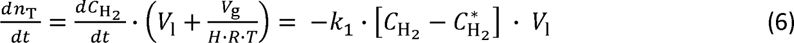

The first-order rate coefficients and their confidence intervals were obtained from the slope of the total mol of H_2_ change per time per volume of liquid culture versus the initial H_2_ concentration minus the H_2_ threshold (rearranging Eq. 6), using the linear regression function of GraphPad 9, which also calculated the 95% confidence interval. Cell-specific first-order rate coefficients 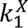 were obtained by dividing *k*_1_ by the cell density or dry weight.

### Determination of first-order kinetic coefficients from time course experiment

To validate the first-order rate coefficients obtained from the initial rate experiment, the kinetic coefficients were compared with first-order coefficients obtained by numerical fitting of our previous experimental H_2_ consumption data (Munoz & Philips, 2023). This study included the three strains used in the initial rate experiment, as well as five additional strains: *S. ovata* (DSM 2663), *Sporomusa sphaeroides* (DSM 2875), *Acetobacterium malicum* (DSM 4132), *Acetobacterium wieringae* (DSM 1911), and *Clostridium autoethanogenum* (DSM 10061), for which also first-order rate coefficients were fitted.

This earlier study (Munoz & Philips, 2023) tested all strains at OD_600_ of 0.2 (similarly as in our initial rate experiment) and used about 25 µM as starting H_2_ concentration, while H_2_ consumption was monitored until the specific threshold was reached. Bottles were first inoculated, after which they were flushed with H_2_ containing gas and heavily shaken for a few minutes to establish a fast equilibrium distribution of H_2_ between the liquid and gas phase (in contrast to the initial rate experiment, in which bottles were inoculated after equilibrium distribution of H_2_).

In addition to these earlier experimental data, the same experiment was repeated for the two *Clostridium* strains at pH 5.9 (instead of pH 7), as this is closer to their optimal pH (Tanner et al., 1993).

The first-order H_2_ consumption coefficient of those eight acetogenic strains was determined by fitting the above ordinary differential equation Eq. (6), on the experimental data, using the starting H_2_ concentration as initial condition. Eq. (6) was numerically integrated by using the *ode15s* algorithm in Matlab. In addition, a non-linear Nelder-Mead simplex algorithm (Lagarias et al., 1998) (*fminsearch*) was used to obtain the fitting curves by minimizing the sum of squares errors (SSE) between the model prediction and the experimental data. Equal weight was applied to all the data points for calculating SSE. The input parameters, *v*_J_, *v*_g_, *H*, *R*, *T*, and 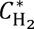 were taken as experimentally determined (Munoz & Philips, 2023). The 95% confidence interval (CI) for each predicted *k*_1_ value was further calculated as described in the **Supplementary Method 2**.

### Evaluation of mass transfer resistance

The calculations described above assume instant equilibrium between the gas and liquid phase. To evaluate the possible influence of mass transfer resistance on our results, we calculated the overall mass transfer coefficient *k*_L_*a* from our orbital shaker settings, based on previously reported mass transfer correlations (Klöckner & Büchs, 2012):

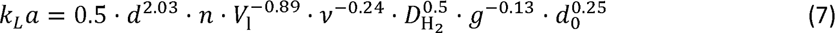

Where *d* is the bottle diameter (0.05 m), *n* the shaking frequency, *V*_l_ is the liquid phase volume, *v* is the kinematic viscosity of the liquid (8×10^−7^ m^2^/s), *D*_H2_ the diffusion coefficient of H_2_ in water (4.5×10^−9^ m^2^/s) (Haynes, 2017), *g* is the acceleration of gravity and *d*_0_ is the shaker diameter (0.025 m). The overall mass transfer coefficient at the side of the gas phase *K*_p_ was calculated using (Cussler, 2009):

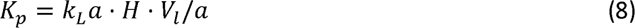

With *a* the area of the interface between the liquid and gas phase.

This mass transfer coefficient was used to evaluate whether H_2_ mass transfer resistance could have impacted the kinetic order type, observed by the initial rate experiment. Hereto, the outcome of the initial rate experiment was simulated using a system of two ODE’s (additional ODE to describe changes in the headspace H_2_ partial pressure *p*_H2_), as derived in **Supplementary Method 3** :

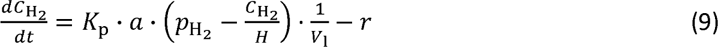

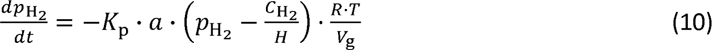

Both equations (9) and (10) include a term for the mass transport of H_2_ from the headspace to the liquid phase, of which the rate depends on the difference in concentration between the two phases and the value of *K*_p_. In addition, Equation (9) accounts for the microbial H_2_ consumption in the liquid phase through the term *r*. We investigated the effect of mass transfer resistance on the kinetic order type observed by the initial rate experiment, assuming either first-order kinetics (*r* given by Equation 4) or Monod kinetics (*r* given by Equation 2). For first-order kinetics, kinetic parameter values as experimentally derived in this study were used (**Table 2**). For Monod kinetics, a *K*_s_ value of 4 µM was used (**Table 1**) and *q*_max_ and its standard deviation were set to match the value of the experimentally observed maximum consumption rate for *S. ovata* (**Figure 1**). The system of ODE’s (Equations 9 and 10) was solved using the *ode15s* algorithm in Matlab, assuming equilibrium between the gas and liquid phase as initial conditions. Five different initial H_2_ concentrations, using approximately the same initial conditions and the same percentage decrease as in the initial rate experiment, were used to model the initial H_2_ consumption rates. These were compared with a simulation assuming instant equilibrium (Eq. 6).

**Figure 1.**
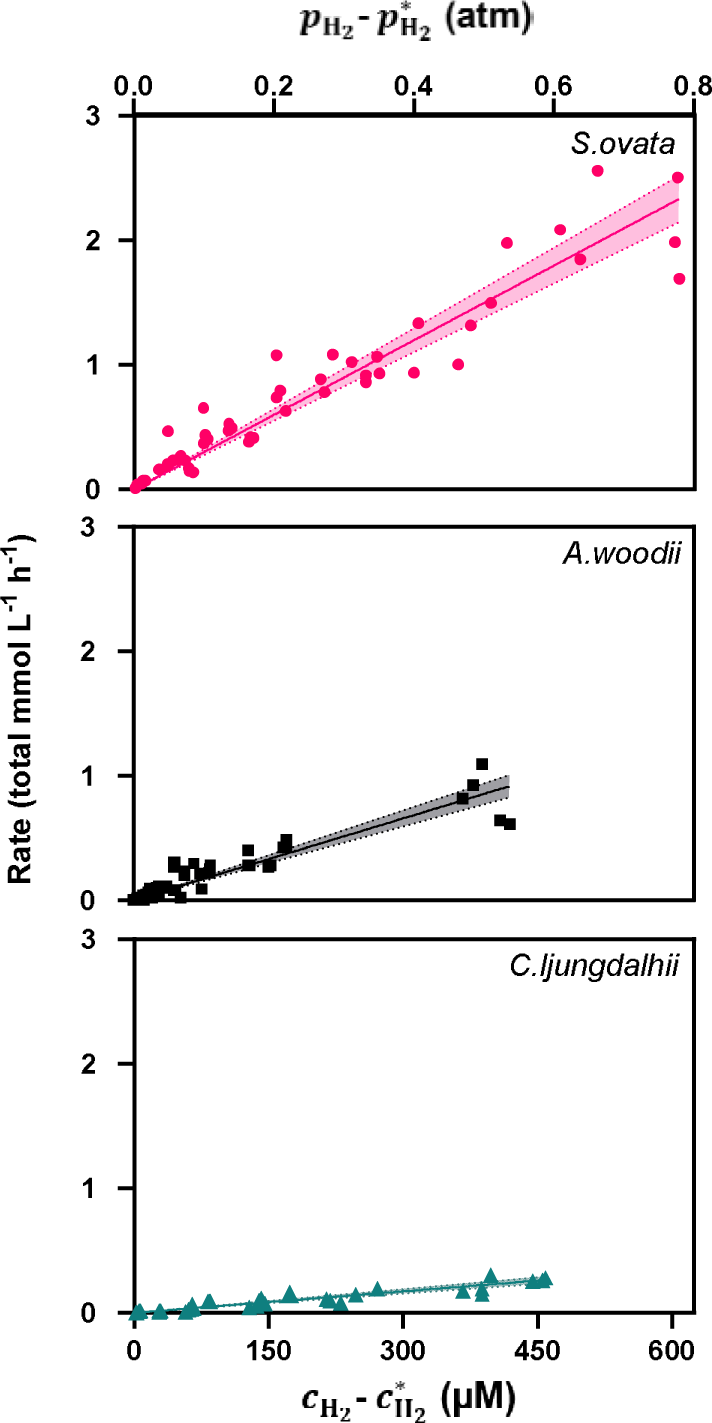
The observed H_2_ consumption rates in function of the initial H_2_ concentration for *S. ovata* (top), *A. woodii* (middle) and *C. ljungdahlii* (bottom), all at pH 7. The observed H_2_ consumption rates (*y*-axis) were expressed as total mmol H per hour per L of liquid phase 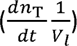 (Eq. 6) to account for the replenishment of H_2_ from the headspace. The H_2_ concentrations (*x*-axis) are corrected by the H_2_ threshold (Munoz & Philips, 2023), thus the origin of the axis corresponds with the respective H_2_ threshold of each strain. All plots use the same scale on the *y*-axis to allow comparison. A plot for *C. ljungdahlii* with adjusted scale is given in **Figure S3**. The linear fit reflects the first-order reaction kinetics. The dotted line around the linear regression represents the 95% confidence interval. The liquid medium is saturated with H₂ at 780 µM in our conditions. Each replicate is shown as a separate dot in the plot.

**Table 2.** First-order rate coefficients (*k*_1_) of H_2_ consumption obtained by the initial rates experiment (IR) (grey) or by fitting the time course experiment (TC) for different acetogenic bacteria. First-order rate coefficients are also reported normalized by the biomass concentration (*k*_1_^X^), expressed as cell density or as gram dry weight (g_DW_) per L. All parameters are at pH 7 unless stated otherwise. The 95% confidence intervals are presented in brackets next to the values. These parameters were obtained under the assumption of instant equilibrium between the gas and liquid phase.

| Organism | Experiment | $k_1$ | $k_1^X$ | $k_1^X$ |
| --- | --- | --- | --- | --- |
| | | $(h^{-1})$ | DW Specific $k_1^X$<br>$(h^{-1} (g_{DW} L^{-1})^{-1})$ | Cell density specific $k_1^X$<br>$\times 10^{-10} (h^{-1} (cell L^{-1})^{-1})$ |
| *S. sphaeroides* | TC | 0.88 (0.79 to 0.97) | 10.63 | 0.06 |
| *C. ljungdahlii* | TC | 0.94 (0.79 to 1.08) | 13.28 | 0.64 |
| *C. ljungdahlii* | IR | 0.58 (0.53 to 0.62) | 8.19 | 0.25 |
| *C. autoethanogenum* pH 5.9 | TC | 1.13 (0.87 to 1.37) | 12.28 | 0.14 |
| *A. woodii* | TC | 1.74 (1.51 to 1.97) | 15.47 | 0.39 |
| *A. woodii* | IR | 2.18 (2.03 to 2.33) | 19.36 | 0.44 |
| *C. ljungdahlii* pH 5.9 | TC | 2.76 (2.65 to 2.87) | 39.02 | 1.11 |
| *S. ovata* 2662 | TC | 3.27 (2.82 to 3.72) | 29.81 | 1.09 |
| *S. ovata* 2662 | IR | 3.84 (3.62 to 4.06) | 34.97 | 1.21 |
| *C. autoethanogenum* | TC | 3.49 (3.03 to 3.95) | 38.19 | 0.44 |
| *S. ovata* 2663 | TC | 3.83 (3.60 to 4.06) | 53.98 | 0.88 |
| *A. malicum* | TC | 2.38 (2.07 to 2.68) | 13.39 | 0.15 |
| *A. wieringae* | TC | 5.79 (5.61 to 5.96) | 60.93 | 0.67 |

Furthermore, we assessed the possible effect of mass transfer resistance on the fitted *k*_1_ values, by repeating the fitting of the first-order kinetic coefficients (as described above), using the system of ODE’s including mass transfer (Equations 9 and 10, using first-order kinetics).

## Results

### Initial consumption rates experiment (IR)

H_2_ consumption rates for at least five different initial H_2_ concentrations were determined in the same conditions for *S. ovata*, *A. woodii* and *C. ljungdahlii*. Biological consumption was confirmed by the faster depletion of H_2_ than in abiotic controls under the same conditions (**Figure S1**). The H_2_ consumption rates were calculated using the initial data points, corresponding to about a 10% change of the initial H_2_ concentration. In all tested conditions, no change of the OD_600_ was expected during the short experimental timeframe, as the amount of consumed H_2_ was insufficient to lead to observable growth, as was confirmed in a parallel experiment (**Figure S2**). In addition, changes in the acetate concentration were insufficient to be detected.

Initially, H_2_ consumption rates were measured up to a dissolved H_2_ concentration of 180 µM (approx. 20% in headspace), since the H_2_ consumption rate was expected to reach a maximum already in the lower µM range, based on previously reported Monod half-saturation coefficients for *A. woodii* and *S. termitida* (Breznak et al., 1988; V. Peters et al., 1998). Plotting the experimentally determined H_2_ consumption rates versus the dissolved H_2_ concentration, however, showed a linear relationship and no indication of a deflection or a maximum rate at higher H_2_ concentrations (**Figure 1, Figure S3**). For this reason, the range of initial concentrations was extended to 450 µM for *A. woodii* and *C. ljungdahlii*, and even to 600 µM for *S. ovata*. Note that the latter concentration is close to the dissolved H_2_ concentration 780 µM at saturation (1 atm of pure H_2_ in headspace at 30 °C) and could not be further increased at the same total pressure and temperature without omitting 20% CO_2_. Even with this extended range of initial H_2_ concentrations, the relationship between the initial H_2_ concentration and the H_2_ consumption rate remained linear, for all three tested strains, without any observable deflection towards a maximum (**Figure 1, Figure S3**). This linear relationship indicates that H_2_ consumption by acetogens follows first-order reaction kinetics, at least at H_2_ concentrations up to saturation (not considering overpressure). The absence of a maximum rate at the highest H_2_ concentrations suggests that H_2_ consumption remains limited by the H_2_ concentration and not by the uptake or conversion capacity of the cells.

It can also be observed from **Figure 1** that *S. ovata* consumed H_2_ faster at the same initial H_2_ concentration than *A. woodii* and *C. ljungdahlii.* The slopes of the plots of **Figure 1** indeed relate to the first-order rate coefficient *k*_1_ (Eq. 6). The first-order rate coefficient is more than six times higher for *S. ovata* than for *C. ljungdahlii* and almost double that for *A. woodii* (**Table 2**). *C. ljungdahlii* has the lowest *k*_1_ among the three strains evaluated in the initial rates experiment, which is likely due to the experimental pH 7, which deviated from its optimum pH 6 (Tanner et al., 1993). Also, when expressed relative to the cell density or dry weight, the cell-specific first-order coefficient (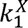) was highest for *S. ovata* (**Table 2**).

### Evaluation of the impact of mass transfer resistance on the kinetic order type

We evaluated whether the H_2_ mass transfer resistance in our system could have impacted the type of kinetics revealed by **Figure 1**. We first estimated the overall mass transfer coefficient from the characteristics from our specific setup and orbital shaker settings using Equations (7) and (8), which gave a *k_L_a* of 29.2 h^−1^(8.1×10^−3^ s^−1^) and a *K*_p_ of 3.6×10^−10^ mol·(m^2^·s·Pa)^−1^ using a shaking speed of 100 rpm.

This value was used to simulate the outcome of an initial rate experiment, under the assumption of first-order or Monod kinetics. Our simulations showed that our initial rate experiment revealed the true kinetic order type, as the shape of the curve was not impacted by the actual mass transfer resistance (**Figure 2**). The figure 2 does show that the slope of the curves and thus the *k*_1_ value (in case of first-order kinetics) is slightly underestimated, when instant equilibrium is assumed under conditions of mass transfer resistance. This is further evaluated below.

**Figure 2.**
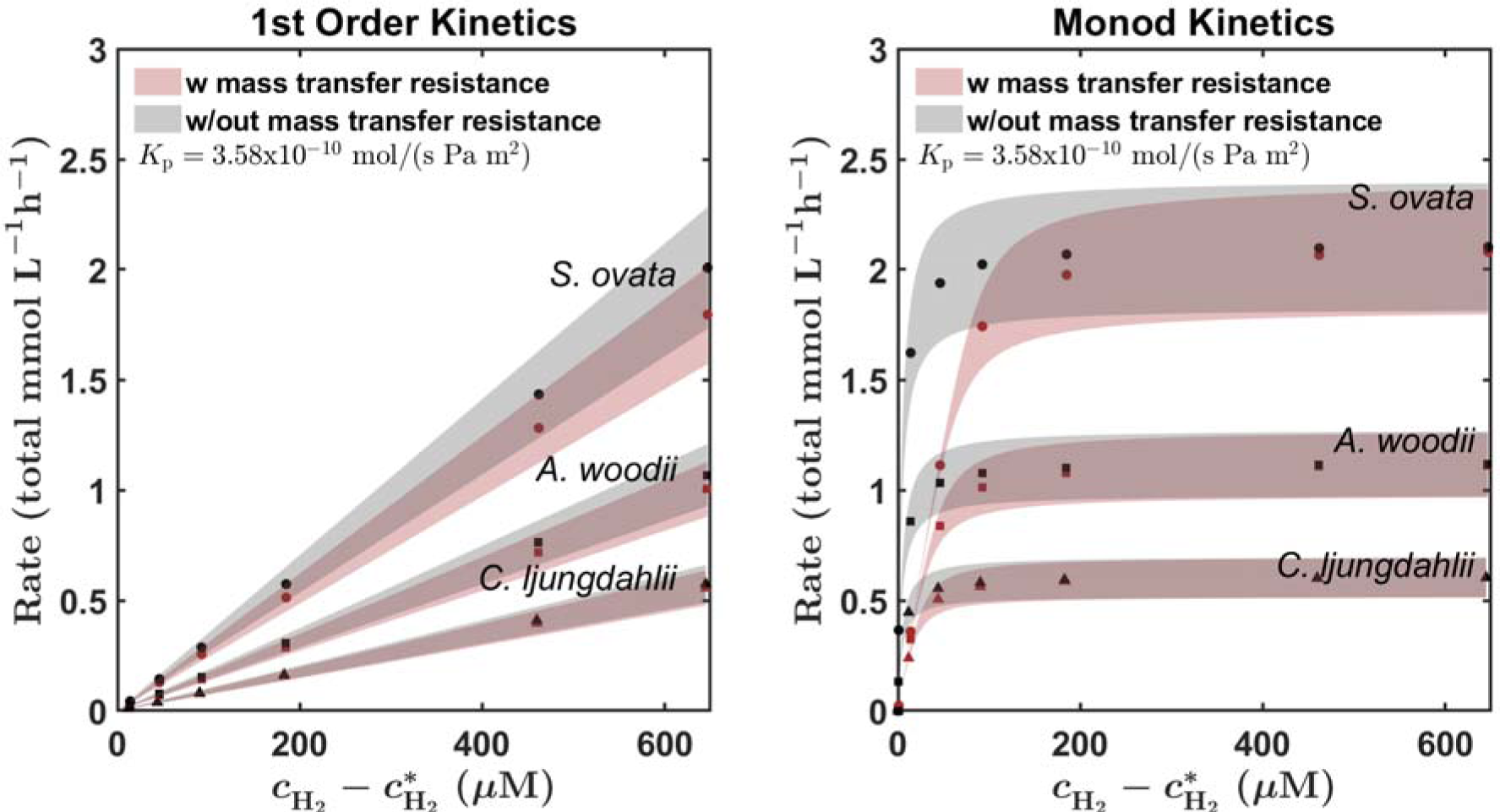
Simulated outcomes of an initial rate experiment under the actual mass transfer resistance (red) or instant equilibrium (black) assuming two types of kinetic orders: left-hand side panel represents first order kinetics, while right-hand side panel reports Monod kinetics. The 95% confidence intervals, evaluated with the experimentally measured standard deviations, are reported as shaded areas. The figure shows that the actual mass transfer resistance in our system did not mask the kinetic order, since the shapes of the simulated curves clearly reflect the type of kinetics under both the actual mass transfer limitations and instant equilibrium.

Our simulations also showed that only at a 50 times lower *K*_p_ (which would correspond to static incubation), mass transfer resistance would be so strong that it would lead to a linear relationship between the consumption rate and the H_2_ concentration, also in case of Monod kinetics (**Figure S4**). In this case, the system would be fully controlled by mass transfer limitations, and there would not be a difference in the observed slope between different strains with different kinetic rates.

### Fitting time course experiment (TC)

To validate the obtained first-order rate coefficients for *S. ovata* 2662, *A. woodii* and *C. ljungdahlii*, the kinetic coefficients were compared with values obtained by fitting Eq. 6 on previous experimental H_2_ consumption time course data (Munoz & Philips, 2023). These independent experiments were performed at the same pH, temperature, medium and agitation, but used only one initial H_2_ concentration (∼25 µM) and followed the consumption of H_2_ until the H_2_ threshold was reached. The fitted curves described the H_2_ depletion reasonably well (**Figure 3**) and the *k*_1_ values obtained using the two experimental approaches are highly comparable (**Figure 4, Table 2**). For *C. ljungdahlii*, the fitted *k*_1_ possibly slightly underestimates the actual value, as for this strain a lag phase of about 1 day was observed in the time course experiment, while all points were included in the fitting. For *S. ovata* 2662 and *A. woodii*, no lag phase was observed. The finding of highly similar *k*_1_ values for these strains using two different experimental approaches thus validates our experiments and results.

**Figure 3.**
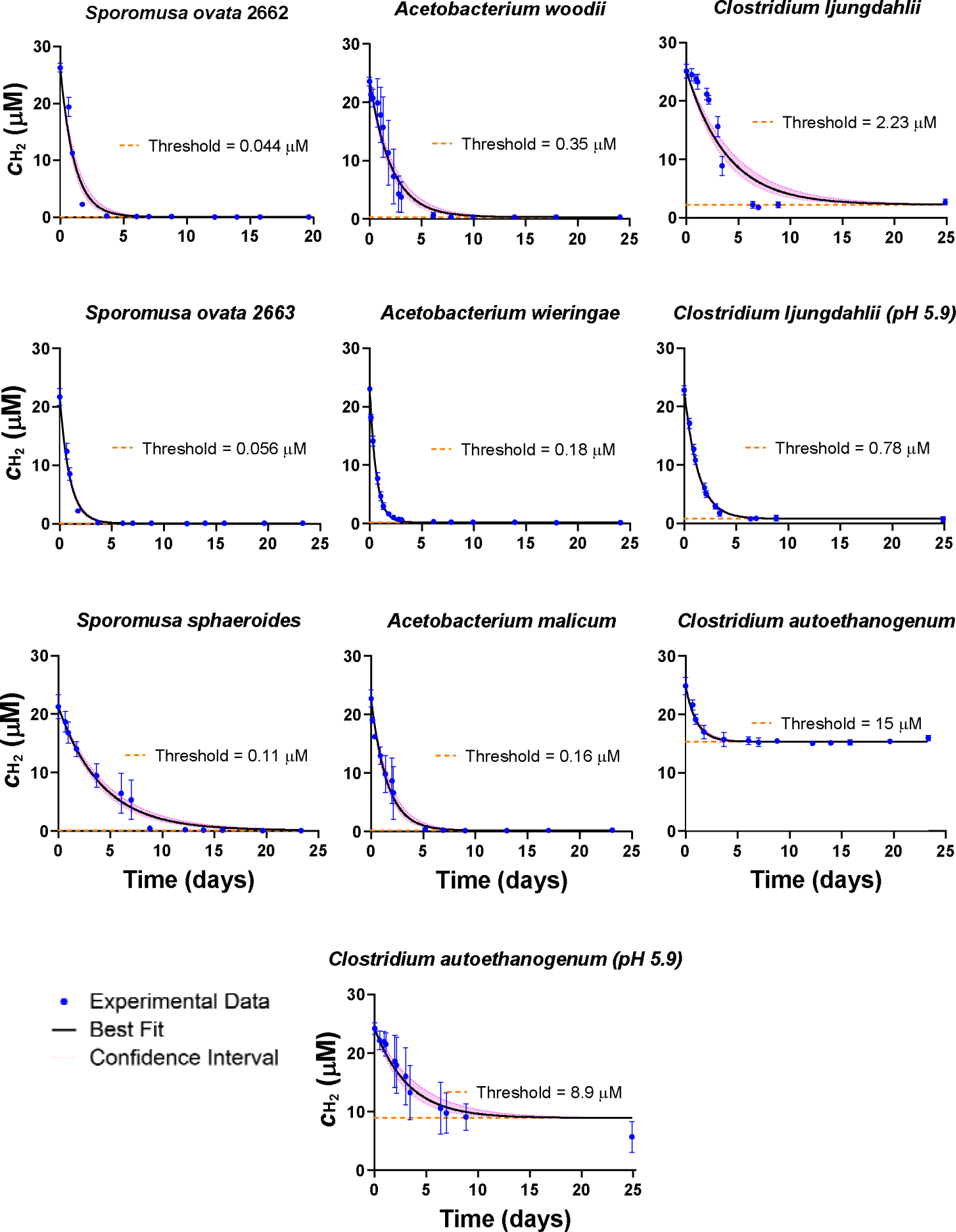
Best-fitting curves on the change of the dissolved H_2_ concentration over time for eight acetogenic strains (Munoz & Philips, 2023) (pH 7), as well as on similar experimental data for *C. autoethanogenum* and *C. ljungdahlii* at pH 5.9. Data points (symbols) represent the mean H_2_ concentration (3 to 5 replicates) with the standard deviation given by the error bars. The best fit is given by the full black lines, while the confidence interval (95%) is aligned by pink dotted lines. The H_2_ threshold is marked by a horizontal orange dotted line.

**Figure 4.**
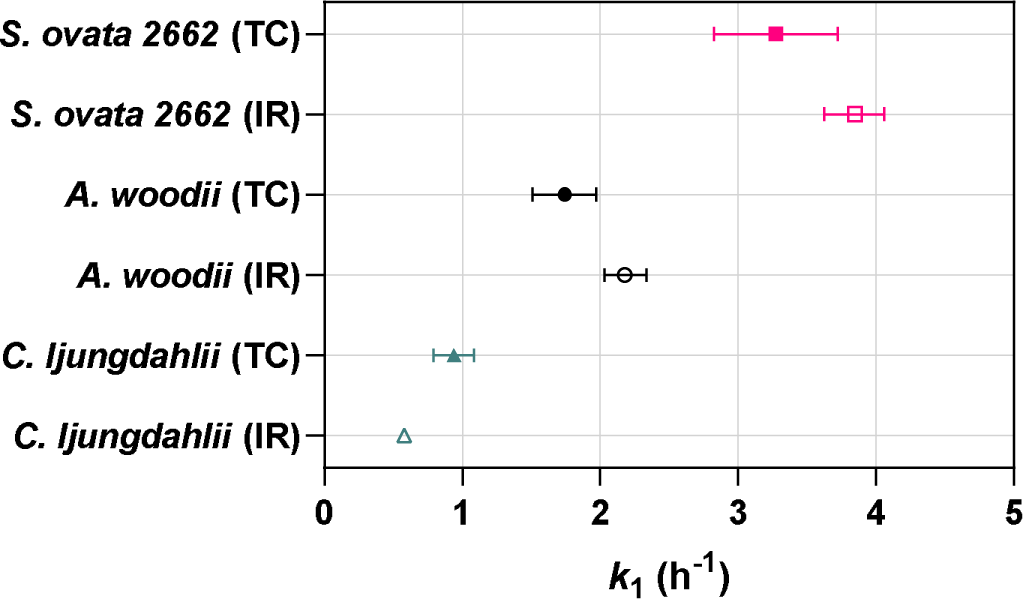
Comparative graph of the numerical values of *k*_1_ obtained from the two experimental approaches: Initial rates experiment (IR) and fitting on the time course experimental data (TC) for *S. ovata* 2662, *A. woodii* and *C. ljungdahlii.* The error bar corresponds to the 95% confidence interval.

In addition, the same fitting method was used to obtain first-order rate coefficients from previous experimental results for five additional acetogenic strains, as well as from new experimental H_2_ depletion data for the *Clostridium* strains at pH 5.9 (**Figure 3**). Biomass-specific first-order rate coefficients *k*_1_^X^ were obtained by normalizing the fitted values by the biomass concentration. We used both dry weights and cell densities to express the biomass concentration, as dry weights are commonly used in literature, but cannot differentiate between viable and dead cells, which is instead feasible with flow cytometry.

Among the tested strains, *A. wieringae* exhibited the highest dry weight specific first-order rate coefficient (**Table 2**, **Figure 5**). Interestingly, both *S. ovata* 2662 and 2663 had significantly higher *k*_1_^X^ values (based on both DW and cell density), compared to *A. woodii*, indicating its higher overall H_2_ consumption capacity under the tested conditions (**Table 2**, **Figure 5, Figure S5**).

**Figure 5.**
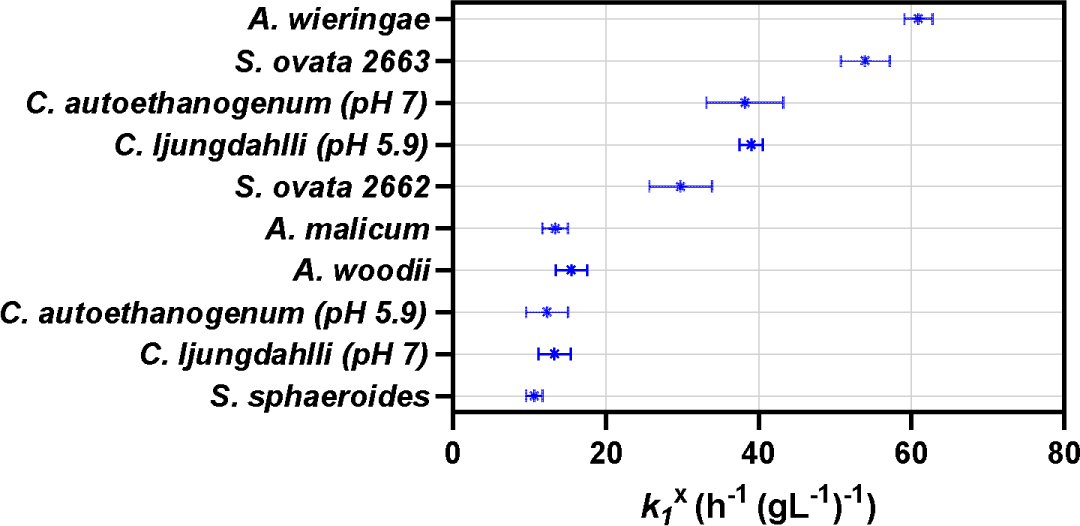
Comparison of the dry weight specific first-order coefficient *k*_1_^X^ obtained by fitting the time course experiment for the acetogenic strains tested. The symbol represents the mean value, and the bars represent the 95% confidence interval.

*C. ljungdahlii* displayed a pH-dependent *k*_1_^X^, with a higher value observed at its more optimal pH 5.9 compared to pH 7 (**Table 2**, **Figure 5, Figure S5**), which aligns with the expected influence of optimal growth conditions on metabolic activity. Notably, *C. ljungdahlii* also demonstrated the highest cell density specific-*k*_1_^X^ at pH 5.9 (**Table 2, Figure S5**) of all the strains tested. *C. autoethanogenum* did not exhibit a higher *k*_1_^X^ at pH 5.9 than at pH 7.0, even though is optimal pH is described to be 5.9 (Abrini et al., 1994). This discrepancy could relate to the narrow concentration difference between the initial H_2_ concentration and the H_2_ threshold for *C. autoethanogenum* at pH 7 (**Figure 3**), which resulted in a wide confidence interval for the fitted *k*_1_ (**Table 2**). Nevertheless, the H_2_ threshold for *C. autoethanogenum* did reflect its more optimal pH at 5.9 (**Figure 3**). Also *C. ljungdahlii* had a lower H_2_ threshold at pH 5.9 than at pH 7.0 (**Figure 3**), while in all conditions the two *Clostridium* strains still had a higher H_2_ threshold, than all other tested strains.

Finally, *S. sphaeroides* displayed the lowest *k*_1_^X^ values (DW and cell density) (**Table 2**, **Figure S5**), but when analyzing the *k*_1_^X^ values expressed by dry weight, no significant differences between *S. sphaeroides* and *A. malicum*, *A. woodii*, *C. autoethanogenum* at pH 5.9, *Clostridium ljungdahlii* were observed (**Table 2**, **Figure 5**).

### Evaluation of the impact of mass transfer resistance on the fitted k_1_ values

Our simulations (**Figure 2**) showed that the assumption of instant equilibrium between the gas and liquid phase could have underestimated the actual *k*_1_ values, which was evaluated further. We evaluated the influence of mass transfer resistance in our system, on the fitted value of the first-order rate coefficient obtained for *A. wieringae*, which is the strain with the highest consumption rate and thus potentially most susceptible to mass transfer resistance. Our analysis revealed that the kinetic coefficient obtained incorporating mass transfer resistance is at the most 32% higher than when assuming gas-liquid equilibrium (**Figure S6**, **Table S2**). For all the other strains, this difference is even lower, due to their lower *k*_1_ (**Table S2**). As this difference is much less than the difference observed between the strains (6-fold difference in *k*_1_ between highest and lowest), as well as the differences that result from other conditions (for instance pH), we conclude that the results of **Table 2** (assuming instant equilibrium) are sufficiently accurate and can be considered as lower limits for the H_2_ consumption rates.

## Discussion

### H_2_ consumption by acetogens follows first-order kinetics and not Monod kinetics up to saturated H_2_ levels

In this work, we found that the H_2_ consumption rates of three different acetogenic strains linearly increased with the dissolved H_2_ concentration, up to almost saturated H_2_ concentrations (maximal tested dissolved H_2_ concentration was 450 to 600 µM) (**Figure 1, Figure S3**). This linear trend shows that H_2_ consumption by the different acetogenic strains followed first-order kinetics. For practical reasons, we did not further increase the H_2_ partial pressure, even though acetogens possibly exhibit a maximum H_2_ consumption rate (e.g. Monod kinetics) at oversaturated H₂ concentrations. Overpressures of H_2_ (>1 atm partial pressure) are relevant for gas fermentation applications, but during microbial electrosynthesis, as well as in most environmental conditions, lower H₂ partial pressures are expected.

Previous studies rather assumed Monod kinetics (Eq. 2) and reported half-saturation coefficients for H_2_ consumption spanning four orders of magnitude (**Table 1**). Previously, Egli (2009) pointed out that determining *K*_s_ is overall difficult, even for soluble substrates, e.g. also for glucose uptake by the model organism *E. coli*, reported *K*_s_ values range over three orders of magnitude. Egli (2009) related this to differences in the experimental approach. In addition, the use of growth rates is problematic, as changes in biomass are hard to measure at low substrate concentrations (Egli, 2009). Also Robinson & Tiedje (1982) pointed out that underestimated H_2_ half-saturation coefficients can result from relating H_2_ consumption to growth. Consequently, the direct determination of the consumption rate in function of the substrate concentration, as in our study, reduces the associated error.

The previous studies reporting half-saturation coefficients for acetogenic H_2_ consumption (**Table 1**) did not report the reliability of their *K*_s_ value. Smith *et al*. (2020) clearly illustrated the importance of reporting the confidence interval of parameter estimates. They found that both Monod kinetics and first-order kinetics fitted batch growth data for the acetogen *Blautia hydrogenotrophica* well, but described that in the case of Monod kinetics, the values of *K*_s_ and *q*_max_ were strongly correlated, which was reflected by an extremely wide confidence interval for *K*_s_. In contrast, they reported a narrow confidence interval for their first-order rate coefficient. In our work, the confidence intervals associated with the first-order kinetic coefficient are also narrow (**Table 2**). In addition, we applied two different experimental approaches (initial rate experiment and fitting time course data), which resulted in highly comparable *k*_1_ values (**Figure 4**). Our *k*_1_ values can thus be considered as highly reliable.

Possibly, H_2_ mass transfer limitations could affect the observed H_2_ consumption kinetic order type (Karadagli et al., 2019; Robinson & Tiedje, 1983). Through simulations (**Figure 2**), we showed that the actual mass transfer resistance in our system could not have masked the true kinetic order type. The mass transfer limitations did affect the estimated *k*_1_ values, but this was maximally 32% for the strain with the highest *k*_1_ (**Figure S6**, **Table S2**).

Besides Smith *et al*. (2020), few other studies previously found indications for first-order H_2_ consumption by acetogens. For instance, a study of *A. woodii* in chemostats to determine its maximal H_2_ uptake rate concluded that a physiological limitation was likely not reached in their experiments (Novak et al., 2021). Also the relatively high *K*_s_ found using several hydrogenase assays (**Table 1**), show that the maximum H_2_ conversion rate was not reached below saturation.

### First-order H_2_ consumption rate coefficients differ up to six times among acetogenic strains

Our results demonstrate that acetogenic strains strongly differ in their first-order H₂ consumption rate coefficient (**Table 2**, **Figure 5**). The highest dry weight specific *k*_1_^X^ coefficient was obtained for *A. wieringae*, which had a six-fold higher consumption rate than *S. sphaeroides,* for which the lowest *k*_1_^X^ value was found. *C. ljungdahlii* at pH 5.9 had one of the highest *k*_1_^X^ expressed per cell density, but an average *k*_1_^X^ expressed per dry weight. This discrepancy likely stems from cell size differences, as *C. ljungdahlii* possibly has larger cells, corresponding to fewer cells per unit of dry weight.

Contrary to what was previously observed for H_2_ thresholds (Munoz & Philips, 2023), *k*_1_^X^ values did not correlate with the genus, indicating that *k*_1_^X^ values might be strain-specific. Likewise, Groher and Weuster-Botz (2016a) described variations in the maximal H_2_ uptake rate and maximal growth rate among various acetogens, including three *Acetobacterium* and two *Sporomusa* species. Differences in growth rates at comparable conditions within the same genus were also observed by Bengelsdorf *et al*. (2016) for four different *Clostridium* strains.

In part, these variations in *k*_1_^X^ are likely due to differences in optimal conditions, as illustrated by the different *k*_1_^X^ values at different pH for the *Clostridium* strains (**Table 2**, **Figure 5**). Besides pH, the H₂ consumption rate of a single acetogenic strain varies with the temperature (Kotsyurbenko et al., 2001) and the medium composition. For example, differences in the concentrations of yeast extract, reductive agents or trace elements (selenium and tungsten) are known to affect acetogenesis rates (An & Kim, 2022; Groher & Weuster-Botz, 2016b; Lanzillo et al., 2024).

In addition to varying optimal conditions, differences in H₂ consumption rates between acetogenic strains could be due to different kinetics of the involved hydrogenases, which are the metalloenzymes catalyzing the redox conversion between H_2_ and protons. The hydrogenases type [NiFe] and the [FeFe] hydrogenases are commonly found in acetogens (Madjarov et al., 2022; J. W. Peters et al., 2015; Schuchmann et al., 2018). These enzymes exhibit complex structural features, including hydrophobic channels for gas diffusion to the active site (Zacarias et al., 2019). This structural conformation can influence the affinity and the sensitivity of the enzyme (Leroux et al., 2008). Indeed, some studies already demonstrated differences in the hydrogenase kinetics of different acetogens (Daniel et al., 1990; Ragsdale & Ljungdahl, 1984; Skidmore et al., 2013).

The *k*_1_^X^ values obtained in this study (**Table 2**) cannot directly be compared with literature values, as previously Monod kinetic parameters were typically reported. To allow comparison, we propose to calculate the biomass-specific H_2_ consumption rate at saturation (multiplying *k*_1_^X^ with the dissolved H_2_ concentration at saturation), which gives rates in the range of 6 to 31 mmol H_2_·g ^−1^·h^−1^ for the acetogens tested in our study (**Table S3**). For other acetogens, Groher & Weuster-Botz (2016a) reported H_2_ consumption rates between 11 to 58 mmol H ·g ^−1^·h^−1^ (**Table S3**), which mostly overlaps with the range found in our study. In addition, Claassens et al., (2019) described biomass-specific H consumption rates of acetogens to range between 14 to 266 mmol H ·g ^−1^·h^−1^, which included thermophilic species. When comparing these values among different studies, it should be taken into account, that these H_2_ consumption rates often were obtained in different conditions (e.g. other medium) and in different experimental setups, which possibly explains why for *A. woodie* the H consumption rate was found to range from 9 to 59 mmol H ·g ^−1^·h^−1^ (**Table S3**). In our study, *A. wieringae* exhibited the highest H_2_ consumption rate under our experimental conditions, followed by *S. ovata* and the two *Clostridium* strains (**Table 2**, **Figure 5**). Numerous studies already explored the capabilities of *C. ljungdahlii* and *C. autoethanogenum* as catalysts in gas conversion technologies (Bardone et al., 2018; Boto et al., 2023; Martínez-Ruano et al., 2023; Valgepea et al., 2022; Zhu et al., 2020), because of their high conversion rates. Maybe less known, but also *A. wieringae* has already been recognized for its high conversion rates on C₁ carbon sources (Arantes et al., 2020; Moreira et al., 2023). In addition, *A. wieringae* was the keystone member in an enriched microbial community leading to high rates of microbial electrosynthesis (Marshall et al., 2017; Mills et al., 2022), making it a highly promising strain for applications.

### Implications

Our finding that H_2_ consumption by acetogens follows first-order kinetics up to almost saturated H_2_ levels, means that the dissolved H_2_ concentration is a major factor limiting acetogenesis in both biotechnological and environmental systems. First-order kinetics entails that the rate of H_2_ conversion per unit biomass is higher at an increased dissolved H_2_ concentration (Eq. 4). This insight is highly relevant to optimize acetogenic biotechnologies. In gas fermentation, increasing the H_2_ partial pressure is often used to increase the H_2_ solubility and thus to overcome mass transfer resistance (Van Hecke et al., 2019). Our results suggest that the high H_2_ partial pressure during gas fermentation also leads to higher consumption rates by acetogens. In contrast, low dissolved H_2_ levels often prevail in microbial electrosynthesis systems, which likely contributes to their limited acetate production rates (Prévoteau et al., 2020).

In addition, the understanding that H_2_ consumption by acetogens follows first-order kinetics is important to understand their role in microbial communities, such as in mixed culture biotechnologies (e.g. anaerobic digestion) or environmental settings. For instance, competition between acetogenic bacteria and methanogenic archaea has been studied by comparing their Monod kinetic parameters for H_2_ consumption (Kotsyurbenko et al., 2001; Tsapekos et al., 2022), but the use of the right kinetic order could lead to more accurate interpretations. Furthermore, to understand acetogenesis in biotechnological and environmental settings, numerical simulations are highly interesting, but the value of models entirely depends on the use of the correct expressions and parameter values governing acetogenic kinetics. Consequently, the insights generated by our study are of high relevance for any research related to acetogenesis.

This work also showed that first-order rate coefficients between acetogens differ up to six times (**Table 2**, **Figure 5**). The *k*_1_^X^ values obtained by this work are of high value for optimal strain selection for the different biotechnological applications. Nowadays, acetogenic strain selection is mostly based on the desired end-product. However, genetic engineering can alter the formed end-product (Bourgade et al., 2021; Moreira et al., 2023; Tremblay & Zhang, 2023), while H_2_ consumption and product formation rates are much harder to engineer. Consequently, it remains important to understand the intrinsic differences in kinetics between the available acetogenic strains to allow optimal strain selection.

Finally, it should be noted that this work only characterized H_2_ consumption rates. This work did not analyze growth nor acetate production rates. Besides the *k*_1_^X^ value of the selected strain, the H_2_ consumption rate also depends on the biomass density in the system (Equation 3), which in turn depends on growth rates, growth yields and cell decay. Acetogens likely also differ in their growth yields, as they have different ATP gains (Munoz & Philips, 2023). Furthermore, product versus biomass yields might vary among acetogens, as well as cell decay rates. Insights into all these aspects are required, before a comprehensive understanding of the product formation and growth kinetics of acetogens becomes possible.

## Author contributions

LM and JP designed the experiments; LM performed the experimental work; SC fitted the *k*_1_ values on our experimental results; MFB performed qPCR; JC performed the simulations investigating mass transfer limitations; LM and LVG analyzed the experimental data; LM and SC constructed the figures; LM and JP wrote the manuscript; SC, FMB, LVG and JC edited the manuscript; JC and JP acquired funding and performed supervision. All authors contributed to the article and approved the submitted version.

## Acknowledgements

This work was financed by a starting grant of Aarhus Universitets Forskningsfond (AUFF), as well as by a Green Transition Project I grant (MES-MODEL, 1127-00048B) from the Independent Research Fund Denmark (DFF).

We thank Lars-Erik Meyer for his advice on how to determine kinetics. We also thank Thomas Dyekjær, Cecilie Nielsen, Gitte Hastrup and Heidi Skov Johansen for their technical and practical support of this project. Miguel Oliveira is acknowledged for his contribution to the Bactobox setup and Ivan Schlembach for the 3D printed adaptor for the manometer.

