## Supplementary Information for "H_2_ consumption by various acetogenic bacteria follows first-order kinetics up to H_2_ saturation"

to

[2] Åbogade 40A,

Aarhus, 8200 Denmark

### Supplementary Method S1

#### DNA isolation and quantitative polymerase chain reaction (qPCR)

For DNA isolation, samples containing *Sporomusa ovata* DSM 2662, *Clostridium ljungdahlii* or *Acetobacterium woodii* were centrifuged at 5000 g during 10 min. The genomic DNA (gDNA) of the cell pellets was extracted using a GeneJET Genomic DNA Purification kit (Thermo Scientific™), following the fabricant's instructions, but using a bead beating step instead of lysozyme. The gDNA was quantified using a Nanodrop spectrophotometer.

The abundance of *Sporomusa ovata*, *Clostridium ljungdahlii* and *Acetobacterium woodii* on the extracted DNA was quantified using quantitative PCR (qPCR). The primers used (Bac534R: 5′-ATTACCGCGGCTGCTGG-3′ and Bac338F: 5′-ACTCCTACGGGAGGCAGCAG-3′) targeted the universal 16s rRNA gene. The qPCR reactions contained 5 µL of iTaq Universal SYBR Green Supermix (2X) (Bio-Rad, Denmark), 1 µL of each primer, 1 µL of gDNA template and 2 µL of nuclease-free water (Thermo Scientific™). The reactions were carried out in a CFX Connect Real-Time PCR Detection System (Bio-Rad). The protocol for thermal cycling consisted in an initial denaturation step at 95 °C for 3 min, followed by 40 cycles consisting of denaturation at 95 °C for 10 s and annealing and extension at 60 °C for 30 s. Samples without DNA were included as negative controls and all the samples were run in duplicates. Melting curves were analyzed to verify the specificity of amplification. Gene counts were corrected for multiple 16S rRNA gene copies at species level for *C. ljungdahlii* (n=9) and *A. woodii* (n=5), and at genus level for *S. ovata* (n=12) (Stoddard et al., 2015).

### Supplementary Method 2

#### Calculation of the 95% Confidence Interval (CI) for non-linear regression using an analytical solution

The ODE with one (number of parameters, *p* = 1) fitting parameter ($k_1$, best fitting being at $\bar{k}$) is as given in the main manuscript by Equation (6):

$$\frac{dC_{\mathrm{H_2}}}{dt} = -k_1 \cdot \left(C_{\mathrm{H_2}} - C^*_{\mathrm{H_2}}\right) \cdot \left(\frac{V_\mathrm{l}'}{V_\mathrm{l}' + V_\mathrm{g}}\right) \tag{S1}$$

Where $k_1$ is the first-order rate coefficient, $C_{\mathrm{H_2}}$ and $C^*_{\mathrm{H_2}}$ are the $H_2$ concentration in the liquid and the $H_2$ threshold concentration, respectively, $V_\mathrm{g}$ is the volume of the gas phase and $V_\mathrm{l}'$ is the 'effective' liquid volume, which relates to the volume of the liquid, $V_l$, the absolute temperature, *T*, and the ideal gas constant *R* and the Henry constant *H* as:

$$V_l' = V_l \cdot H \cdot R \cdot T \tag{S2}$$

Equation S1 must be solved together with the initial condition, in which $C^0_{\mathrm{H_2}}$ is the initial $H_2$ concentration:

$$C_{\mathrm{H_2}}(t = 0) = C^0_{\mathrm{H_2}} \tag{S3}$$

It should be noted that these equations assume instant equilibrium between the liquid and gas phase (no mass transport resistance) and that the measurable parameter was the $H_2$ partial pressure $p_{\mathrm{H_2}}$, which allows to calculate the $H_2$ concentration using the Henry's constant *H*.

By integrating the ODE we obtained the following analytical solution:

$$C_{\mathrm{H_2}}(t) = C^*_{\mathrm{H_2}} + \exp\left(-\frac{V_\mathrm{l}' \cdot k_1}{V_\mathrm{g} + V_\mathrm{l}'} \cdot t\right) \cdot \left(C^0_{\mathrm{H_2}} - C^*_{\mathrm{H_2}}\right) \tag{S4}$$

Next, we defined the partial derivative of $C_{\mathrm{H_2}}$ with respect to the fitting parameter $k_1$ as:

$$\frac{\partial C_{\mathrm{H_2}}}{\partial k_1} = \frac{V_\mathrm{l}'}{V_\mathrm{g} + V_\mathrm{l}'} \cdot \exp\left(-\frac{V_\mathrm{l}' \cdot k_1}{V_\mathrm{g} + V_\mathrm{l}'} \cdot t\right) \cdot \left(C^*_{\mathrm{H_2}} - C^0_{\mathrm{H_2}}\right) \cdot t \tag{S5}$$

The analytical Jacobian matrix, $\boldsymbol{J}$, (Bard, 1974) (in this case a vector of *n* elements corresponding to the number of experimental observations – including the replicates) calculated at the optimal solution found with the simplex algorithm $\overline{k_1}$ at each experimental time reads as:

$$\boldsymbol{J} = \begin{bmatrix} \left.\frac{\partial C_{\mathrm{H_2}}(k_1)}{\partial k_1}\right|_{\overline{k_1},t_1} \\ \vdots \\ \left.\frac{\partial C_{\mathrm{H_2}}(k_1)}{\partial k_1}\right|_{\overline{k_1},t_n} \end{bmatrix} \tag{S6}$$

The standard error, *se*, at $k_1 = \overline{k_1}$ is computed as (Bard, 1974):

$$se(k_1) = [(\boldsymbol{J}^T \cdot \boldsymbol{J})^{-1} \cdot \hat{\sigma}^2]^{\frac{1}{2}} \tag{S7}$$

With the variance estimate being the sum of squares due to error (SSE) divided by $n - p$ (observation – parameters):

$$\hat{\sigma}^2 = \frac{1}{n-p}\sum_{i=1}^{n}\left(C_{\mathrm{H_2}}(t_i, \overline{k_1}) - C_{\mathrm{H_2}}^{\exp}(t_i)\right)^2 \tag{S8}$$

In Eq. S8, $C_{\mathrm{H_2}}^{\exp}(t_i)$ is the experimentally observed $H_2$ concentration at time, $t_i$. Finally, the 95% CI was calculated from the *t*-distribution as:

$$95\%\ CI = \overline{k_1} \pm t(0.975, n-p) \cdot se(k_1) \tag{S9}$$

### Supplementary Method S3

**Derivation of the system of ODE's used to include $H_2$ mass transfer resistance**

To account for the mass transfer resistance, the starting point is $H_2$ molar balance in the liquid phase:

$$\frac{dn_{\mathrm{H_2,liq}}}{dt} = \dot{n}^{\mathrm{in}}_{\mathrm{H_2,liq}} - \dot{n}^{\mathrm{out}}_{\mathrm{H_2,liq}} + \dot{n}^{\mathrm{g}}_{\mathrm{H_2,liq}}$$

$$= K_{\mathrm{p}} \cdot a \cdot \left(p_{\mathrm{H_2}} - \frac{C_{\mathrm{H_2}}}{H}\right) - k_1 \cdot \left(C_{\mathrm{H_2}} - C^*_{\mathrm{H_2}}\right) \cdot V_{\mathrm{l}} \qquad \text{(S10)}$$

Together with the one for gas phase:

$$\frac{dn_{\mathrm{H_2,gas}}}{dt} = -K_{\mathrm{p}} \cdot a \cdot \left(p_{\mathrm{H_2}} - \frac{C_{\mathrm{H_2}}}{H}\right) \qquad \text{(S11)}$$

In the equations above, $n_{\mathrm{H_2,liq}}$ and $n_{\mathrm{H_2,gas}}$ represent the numbers of moles of $H_2$ in the liquid and gas phase respectively, while $\dot{n}^{\mathrm{in}}_{\mathrm{H_2,liq}}$, $\dot{n}^{\mathrm{out}}_{\mathrm{H_2,liq}}$ , $\dot{n}^{\mathrm{g}}_{\mathrm{H_2,liq}}$ are the molar fluxes of $H_2$ entering, exiting, and being consumed in the liquid phase, respectively. The parameter $k_1$ is the first order microbial $H_2$ consumption kinetic coefficient and $K_{\mathrm{p}}$ is the overall mass transfer coefficient at the gas side. Finally, $a$ is the area of the gas liquid interface.

Rearranging Equations (S10) and (S11) in terms of the $H_2$ concentration in the liquid phase and the $H_2$ partial pressure of the gas phase (the experimentally accessible measurement), the system of ODE's becomes:

$$\frac{dC_{\mathrm{H_2}}}{dt} = K_{\mathrm{p}} \cdot a \cdot \left(p_{\mathrm{H_2}} - \frac{C_{\mathrm{H_2}}}{H}\right) \cdot \frac{1}{V_{\mathrm{l}}} - k_1 \cdot \left(C_{\mathrm{H_2}} - C^*_{\mathrm{H_2}}\right) \qquad \text{(S12)}$$

$$\frac{dp_{\mathrm{H_2}}}{dt} = -K_{\mathrm{p}} \cdot a \cdot \left(p_{\mathrm{H_2}} - \frac{C_{\mathrm{H_2}}}{H}\right) \cdot \frac{R \cdot T}{V_{\mathrm{g}}} \qquad \text{(S13)}$$

These equations are also reported in the main text as Equations (9) and (10).
This ODE system must be coupled with the initial conditions for the $H_2$ concentration and partial pressure:

$$C_{\mathrm{H_2}}(t=0) = C^0_{\mathrm{H_2}} \qquad \text{(S14)}$$

$$p_{\mathrm{H_2}}(t=0) = \frac{C^0_{\mathrm{H_2}}}{H} \qquad \text{(S15)}$$

The first initial condition is identical to Equation (S3), while the second initial conditions states that the gas and liquid phase are in equilibrium at the beginning of the experiment.

If the resistance in the gas phase is negligible with respect to the resistance in the liquid phase (as is often the case), the overall mass transfer coefficient for the gas phase can be replaced with the mass transfer coefficient in the liquid phase ($k_L a$) and Equations (S12) and (S13) can be rewritten as:

$$\frac{dC_{\mathrm{H_2}}}{dt} = k_L a \cdot \left(C_{\mathrm{H_2,eq}} - C_{\mathrm{H_2}}\right) - k_1 \cdot \left(C_{\mathrm{H_2}} - C^*_{\mathrm{H_2}}\right) \tag{S12'}$$

$$\frac{dp_{\mathrm{H_2}}}{dt} = -k_L a \cdot \left(C_{\mathrm{H_2,eq}} - C_{\mathrm{H_2}}\right) \cdot \frac{R \cdot T \cdot V_l}{V_{\mathrm{g}}} \tag{S13'}$$

Where $C_{\mathrm{H_2,eq}} = p_{\mathrm{H_2}} \cdot H$ is the virtual $H_2$ equilibrium concentration in the liquid phase. Equations (S12') and (S13') can be then solved together with the initial conditions in Equations (S14) and (S15).

### Supplementary Figures

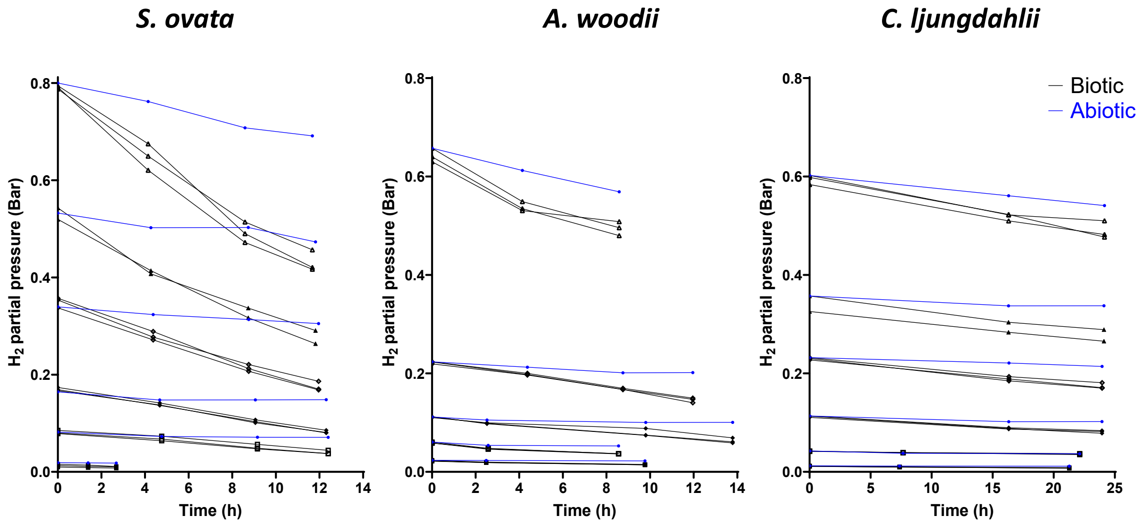


**Figure S1.** Change of the $H_2$ partial pressure in the gas phase over time by *Sporomusa ovata* (left), *Acetobacterium woodii* (middle) and *Clostridium ljungdahlii* (right) (black lines) in comparison to the abiotic controls (blue line) at different initial $H_2$ concentrations (y-axis). Time zero in these graphs corresponds to the initial sampling point. The $H_2$ consumption rate (**Figure 1, main text**) was calculated using the initial sampling points (about 10% change of the $H_2$ concentration). Sampling at additional time points was performed, as it was unknown when the 10% change of the $H_2$ concentration would be reached.

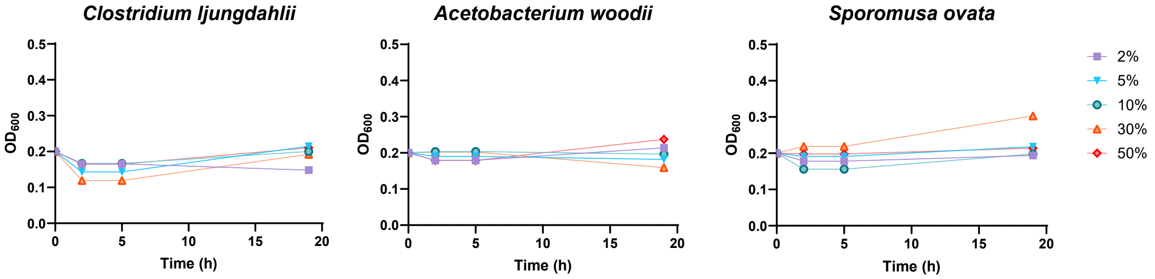


**Figure S2.** Change of the optical density ($OD_{600}$) over time, as measured in a parallel experiment with the same initial $H_2$ concentrations (different colors and symbols) and the same strains as in the initial rate experiment. Over the course of 20 h, the $OD_{600}$ remained stable, allowing the assumptions of constant biomass concentrations in our calculations.

Given the limited amount of $H_2$ available and the relatively high initial cell density, the increase in biomass was anyway expected to be small. In the experiment with the highest $H_2$ concentration (50% at a total pressure of 1.2 bar), the expected biomass increase from consuming 10% of the available $H_2$ is about 0.011 $g_{DW}$/L, using a growth yield of 2.8 $g_{DW}$ of biomass per mole of $H_2$ consumed (own experimental results for *S. ovata*). This is a small increase (15%) in comparison to the initial biomass concentration of 0.071 $g_{DW}$/L (for *Sporomusa ovata*) and equivalent to an $OD_{600}$ increase of about 0.03, while the starting OD was 0.20.

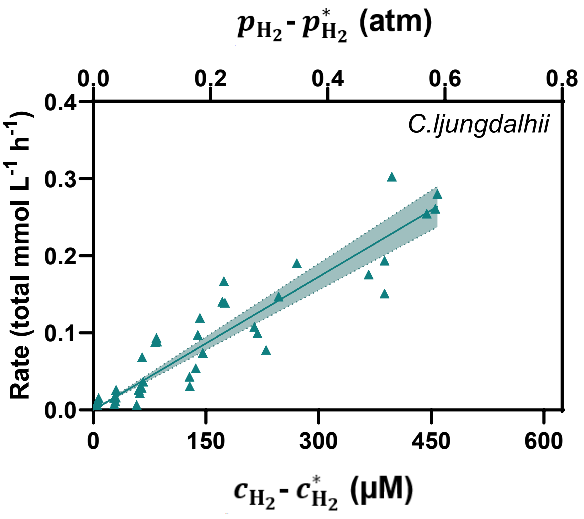


**Figure S3:** The observed $H_2$ consumption rates in function of the initial $H_2$ concentration for *C. ljungdahlii* at pH 7. This plot is the same as in **Figure 1** of main text, but the scale on the Y-axis is adjusted to better visualize the linear relationship between the rate and the initial $H_2$ concentration. This linear relationship reflects first-order reaction kinetics.

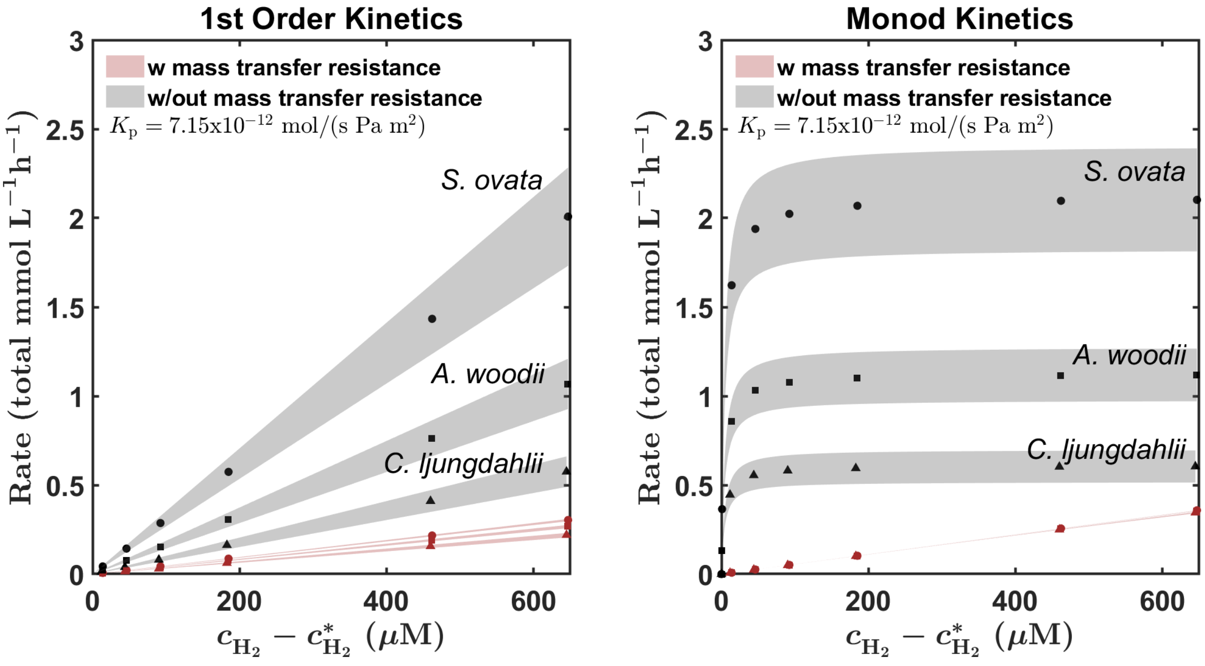


**Figure S4.** Simulated outcome of an initial rate experiment assuming a fifty times lower $K_p$ than calculated for our experimental conditions. These simulations compare the outcome (for *S. ovata, A. woodii,* and *C. ljungdahlii*) under mass transfer resistance (red) or instant equilibrium (black) assuming two types of kinetic orders: left-hand side panel represents first order kinetics, while right-hand side panel reports Monod kinetics.

These simulations used **Equation 6 (main text)** to describe instant equilibrium, while the system of ODEs (**Equations 9 and 10, main text**) was used to model the inclusion of mass transfer resistance. The 95% confidence intervals, evaluated using the experimentally measured standard deviations, are reported as shaded areas.

The figure shows that at a fifty-times lower $K_p$ our experimental system would mask the kinetic order, since a linear relationship would be observed, also in the case of Monod kinetics. It should be noted that under this strong mass transfer resistance, no differences in the slope between the different microbes would be expected.

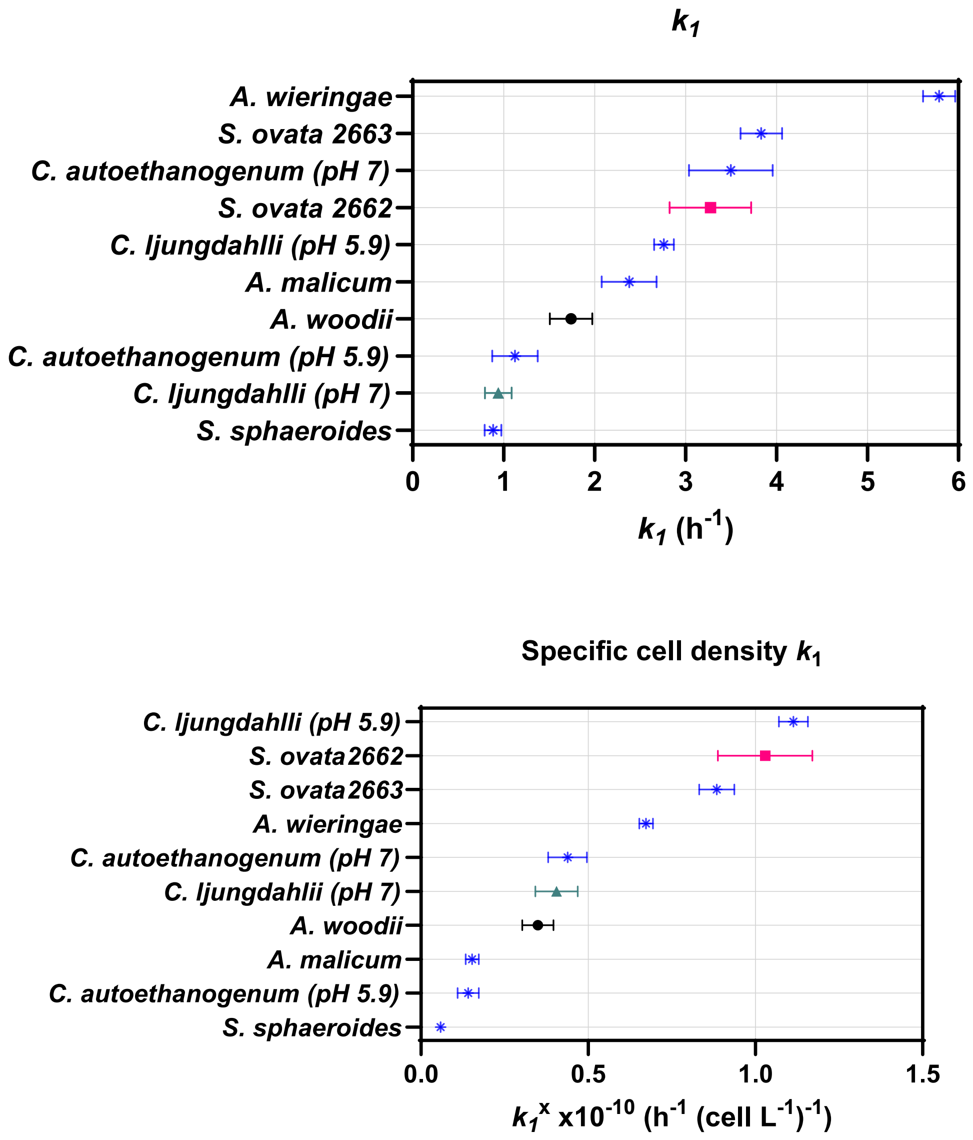


**Figure S5.** Comparative graphs of the highest $k_1$ (top) and cell density specific $k_1^X$ (bottom) obtained by the fitting the time course experiment for all the acetogenic strain tested.

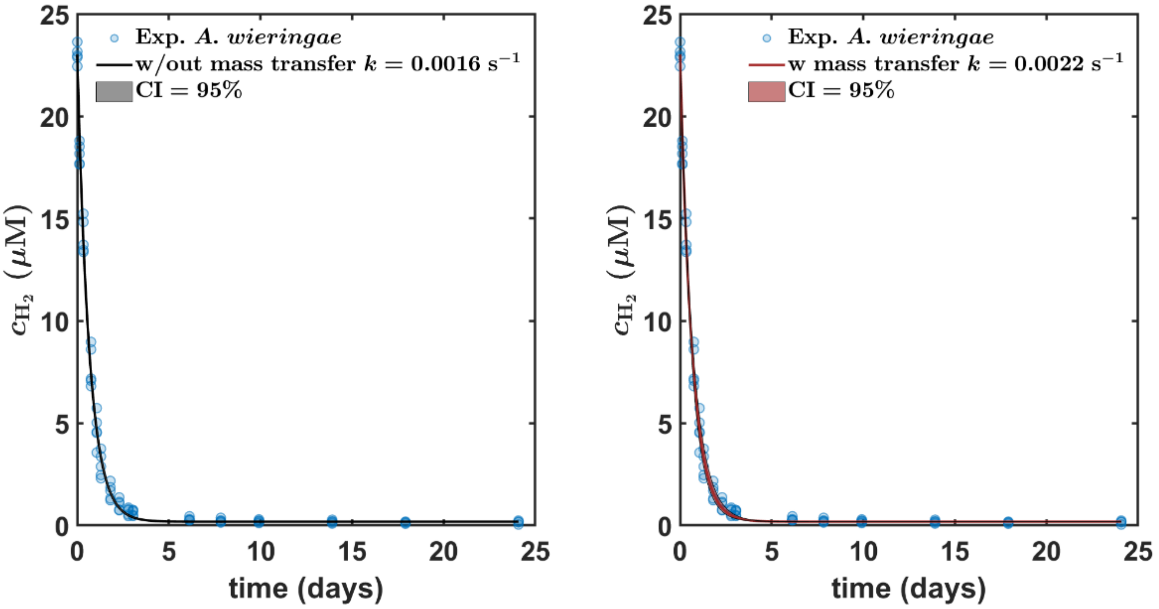


**Figure S6.** Comparison of the fitting in absence (left-hand side panel) and presence (right-hand side panel) of mass transfer resistance. Here we report the fitting for *A. wieringae*, since this species was found to have the highest $H_2$ consumption first-order rate coefficient (**Table 2, main text**), thus the one most affected by mass transfer resistance. These simulations used **Equation 6** (main text) to describe instant equilibrium, while the system of ODE's (**Equations 9 and 10**) (main text) and the $K_p$ calculated for our system (**Equations 7 and 8,** main text) was used to model the inclusion of mass transfer resistance.

The figure shows that both models fit the kinetics very accurately, but that the fitted $k_1$ value with mass transfer resistance (0.0020 s$^{-1}$) was higher than with the assumption of instant equilibrium (0.0015 s$^{-1}$), which is a difference of 34% (**Table S2**). This illustrates that the assumption of instant equilibrium slightly underestimates the actual first order rate coefficients. For all other strains, even lower differences were found, because of their lower $k_1$ values (**Table S2**).

### Supplementary Tables

**Table S1.** The correlation factors from units of OD to dry weight ($g_{DW}$/L) and cell density (cells/mL) for different acetogenic bacteria. Methods are described in the main text.

| Organism | Dry weight $g_{DW}$/L per $OD_{600}$ unit | $R^2$ | Cell density Cells/mL per $OD_{600}$ unit | $R^2$ |
|---|---|---|---|---|
| *Sporomusa ovata* 2663 | 0.355 | 0.988 | 2.17 x$10^8$ | 0.981 |
| *Sporomusa ovata* 2662 | 0.549 | 0.976 | 1.59 x$10^8$ | 0.987 |
| *Acetobacterium woodii* | 0.563 | 0.952 | 2.49 x$10^8$ | 0.997 |
| *Clostridium ljungdahlii* | 0.354 | 0.998 | 1.16 x$10^8$ | 0.971 |
| *Acetobacterium wieringae* | 0.475 | 0.998 | 4.30 x$10^8$ | 0.958 |
| *Acetobacterium malicum* | 0.889 | 0.993 | 7.77 x$10^8$ | 0.994 |
| *Acetobacterium sphaeriodes* | 0.416 | 0.988 | 4.64 x$10^8$ | 0.998 |
| *Clostridium autoethanogenum* | 0.458 | 0.989 | 3.99 x$10^8$ | 0.999 |

**Table S2.** Best fitting and standard deviation (as 95% of the confidence intervals) of the first order kinetic coefficient $k_1$ obtained from our experimental data, assuming gas-liquid equilibrium, or incorporating the actual mass transfer resistance. The difference in the value for $k_1$ is at the most 34% (*A. wieringae*).

| | Gas-liquid equilibrium | | With mass transfer | | % Dif |
|---|---|---|---|---|---|
| | $k_1$ 1/h | SD 1/h | $k_1$ 1/h | SD 1/h | |
| *Sporomusa ovata* 2662 | 3.38 | 0.43 | 3.97 | 0.72 | 17% |
| *Acetobacterium wieringae* | 5.85 | 0.17 | 7.86 | 0.88 | 34% |
| *Clostridium ljungdahlii* pH = 7 | 0.92 | 0.14 | 0.96 | 0.16 | 4% |
| *Acetobacterium malicum* | 2.36 | 0.29 | 2.64 | 0.33 | 12% |
| *Acetobacterium woodii* | 1.71 | 0.23 | 1.85 | 0.27 | 8% |
| *Clostridium ljungdahlii* pH = 5.9 | 2.76 | 0.10 | 3.14 | 0.22 | 14% |
| *Sporomusa sphaeroides* | 0.89 | 0.09 | 0.92 | 0.10 | 4% |
| *Sporomusa ovata* 2663 | 3.84 | 0.21 | 4.61 | 0.44 | 20% |
| *Clostridium autoethanogenum* pH = 7.0 | 3.28 | 0.38 | 3.83 | 0.56 | 17% |
| *Clostridium autoethanogenum* pH = 5.9 | 1.09 | 0.24 | 1.15 | 0.27 | 5% |

**Table S3**: $H_2$ consumption rates close to saturation calculated from this study in comparison to $H_2$ consumption rates reported by earlier studies. All $H_2$ consumption rates were normalized per biomass concentration. $H_2$ consumption rates close to saturation were calculated from our results by multiplying $k_1^X$ (**Table 2,** taken for time course experiment) with the dissolved $H_2$ concentration at a 60% $H_2$ headspace at 1.2 bar (560 µM). Comparable values for the $H_2$ consumption rate were obtained from the data of Groher & Weuster-Botz, (2016), by dividing the reported maximum $H_2$ uptake rate with the reported maximum biomass concentration. Alternatively, a $H_2$ consumption rate in the same unit can be obtained, as $q_{max}$ (mmol $H_2 \cdot g_{DW}^{-1} \cdot h^{-1}$) or by dividing maximum growth rates $\mu_{max}$ ($h^{-1}$) with the growth yield ($g_{DW}$ /mmol $H_2$).

| **Acetogen** | **$H_2$ consumption rate close to saturation (mmol $H_2 \times g_{DW}^{-1} \times h^{-1}$)** | **Source** |
|---|---|---|
| *Acetobacterium fimetarium* | 11.36 | (Groher & Weuster-Botz, 2016) |
| *Acetobacterium malicum* | 7.51 | This study |
| *Acetobacterium wieringae* | 34.21 | This study |
| *Acetobacterium wieringae* | 31.25 | (Groher & Weuster-Botz, 2016) |
| *Acetobacterium woodii* | 8.68 | This study |
| *Acetobacterium woodii* | 58.86 | (Groher & Weuster-Botz, 2016) |
| *Acetobacterium woodii* | 14.18 | (Peters et al., 1998) |
| *Acetonema longum* | 19.25 | (Kane & Breznak, 1991) |
| *Blautia hydrogenotrophica* | 25.60 | (Groher & Weuster-Botz, 2016) |
| *Clostridium autoethanogenum (*pH 5.9) | 6.89 | This study |
| *Clostridium ljungdahlii* (pH 5.9) | 21.91 | This study |
| *Clostridium magnum* | 46.29 | (Groher & Weuster-Botz, 2016) |
| *Eubacterium aggregans* | 40.69 | (Groher & Weuster-Botz, 2016) |
| *Sporomusa acidovorans* | 46.32 | (Groher & Weuster-Botz, 2016) |
| *Sporomusa ovata* 2662 | 16.74 | This study |
| *Sporomusa ovata* 2662 | 46.32 | (Groher & Weuster-Botz, 2016) |
| *Sporomusa ovata* 2663 | 30.31 | This study |
| *Sporomusa sphaeroides* | 5.98 | This study |
